# Microglial Lag3 Drives α-Synuclein–induced Neurotoxic Activated (A1) Astrocytes and Neurodegeneration

**DOI:** 10.64898/2026.01.06.697996

**Authors:** Xiuli Yang, Ramhari Kumbhar, Bo Am Seo, Shinwon Ha, Jared T. Hinkle, Ning Wang, Shuya Li, Lili Niu, Haiqing Liu, Nicolas Stanciu, Rong Chen, Yasuyoshi Kimura, Enquan Xu, Fangyi Cheng, Sung-Ung Kang, Mingyao Ying, Han Seok Ko, Valina L. Dawson, Ted Dawson, Xiaobo Mao

**Author notes:** Corresponding authors (XM); (VLD); (TMD). Department of Convergence Medicine, Department of Biochemistry, Department of Global Medical Science, Organelle Medicine Research Center, Yonsei University Wonju College of Medicine, Wonju-si, Gangwon-do 26426, Republic of Korea; Natural Product Research Center, Korea Institute of Science and Technology, Gangneung-si, Gangwon-do 25451, Republic of Korea (Bo Am Seo).

## Abstract

**Background:** Neuroinflammation and pathologic α-synuclein (α-syn) aggregation cooperate to drive dopaminergic neurodegeneration in Parkinson’s disease, but the glial receptors that couple extracellular α-syn to inflammatory cascades remain incompletely defined. Microglia express higher levels of lymphocyte activation gene 3 (Lag3) than neurons, yet the contribution of microglial Lag3 to α-syn recognition, glial crosstalk, and neurodegeneration is unknown.

**Methods:** Biochemical binding assays, live-cell imaging, cytokine profiling, and neuron–microglia–astrocyte co-culture paradigms were used to define Lag3-dependent α-syn preformed fibril (PFF) binding, uptake, and microglial activation. To interrogate in vivo function, microglia-specific Lag3 conditional knockout mice (Lag3^L/L^-Cx3cr1^CreER^) and littermate controls received unilateral intrastriatal α-syn PFF injections, followed by histological, biochemical, and behavioral assessments of α-syn pathology, gliosis, nigrostriatal integrity, and motor performance.

**Results:** α-syn PFFs bound microglial Lag3 with high specificity and nanomolar affinity and required Lag3 for efficient fibril internalization and induction of proinflammatory cytokines. Microglial Lag3 deficiency markedly blunted α-syn PFF–evoked microglial activation, prevented cytokine-driven conversion of astrocytes into neurotoxic reactive A1 astrocytes, and abolished astrocyte-dependent neuronal death in vitro. In vivo, microglia-specific Lag3 deletion reduced cortical, striatal, and substantia nigra pS129 α-syn pathology, suppressed microgliosis and A1 astrocyte induction, preserved substantia nigra dopaminergic neurons and striatal dopamine transporter/tyrosine hydroxylase expression, and ameliorated α-syn PFF–induced motor deficits.

**Conclusions:** This study identifies microglial Lag3 as a key receptor linking extracellular α-syn PFF recognition to inflammatory amplification, neurotoxic reactive A1 astrocyte conversion, and dopaminergic neurodegeneration. Together with prior work on neuronal Lag3, these findings support a cell-type–specific dual-axis model in which neuronal Lag3 mediates α-syn propagation while microglial Lag3 drives glia-dependent neurotoxicity, positioning Lag3 as a promising precision therapeutic target in α-synucleinopathies.

## Introduction

Parkinson’s disease (PD) is characterized by the progressive misfolding, aggregation, and propagation of α-synuclein (α-syn), along with neuroinflammation and the selective loss of dopaminergic neurons [1–6]. Mounting evidence indicates that α-syn preformed fibrils (PFFs) spread between neurons and glial cells in a prion-like manner [7–9]; however, the cellular receptors that initiate this process remain incompletely defined. Among the proposed candidates, the immune checkpoint receptor lymphocyte activation gene 3 (Lag3) plays a role in the uptake of pathologic α-syn in neurons [10–13].

Although early studies suggested that neuronal Lag3 serves as a high-affinity receptor for α-syn PFFs, facilitating their binding, uptake, and neuron-to-neuron propagation [12], one report contended that Lag3 is not present in neurons and does not facilitate the propagation of pS129-positive α-synuclein pathology [14]. This conclusion contrasts with a body of work reporting detectable neuronal Lag3 expression in multiple species including humans [11–13, 15–22], implicating the receptor in α-synucleinopathy-related pathology [11–13, 23–26]. Analyses of publicly available datasets indicate measurable, albeit low, neuronal Lag3 mRNA levels across several species [15, 17–22]. In line with these observations, single-cell sequencing and complementary data from immunohistochemistry, immunoblotting, and RNAscope in wild-type versus Lag3 knockout mice, together with results from a reporter line harboring a YFP tag in the Lag3 locus, collectively substantiate that neurons do express Lag3 [11]. In addition, loss of Lag3 markedly reduces phosphorylated serine 129 (pS129)-positive α-syn pathology and alleviates behavioral deficits in transgenic mice overexpressing human A53T α-syn [23]. Mice with selective deletion of Lag3 in neurons also exhibit reduced pS129 α-syn pathology, reduced loss of DA neurons, and behavioral deficits after intrastriatal α-syn PFFs injection [25]. These results indicate that neuronal Lag3, despite low abundance, is functionally relevant in mediating pathogenic α-syn transmission. Consistent findings from independent studies show that Lag3 deficiency in microvascular endothelial cells attenuates α-syn spread, neuronal loss, and related behavioral abnormalities [26]. When considered alongside earlier in vitro and in vivo evidence implicating Lag3 in the transmission of pathological α-syn, these results support a central role for Lag3 in mediating the dissemination of toxic α-syn species. Moreover, preclinical data indicate that Lag3 knockout mice are protected from α-syn PFF–induced dopaminergic neurodegeneration and from gut-to-brain α-syn transmission [24].

Although neuronal Lag3 expression is low, microglia consistently express higher baseline levels of Lag3 [27–29]. Microglia are also among the earliest responders to extracellular pathologic α-syn [2, 30, 31]. Microglia can initiate potent inflammatory responses that convert astrocytes into neurotoxic reactive (A1) astrocytes, a process now recognized as a major contributor to neuronal vulnerability in neurodegeneration [32, 33]. Despite the higher levels of Lag3 in microglia, the role of microglial Lag3 in sensing pathologic α-syn and in driving downstream glial toxicity has remained unexplored.

In this study, we identify microglial Lag3 as a key receptor for extracellular pathologic α-syn and demonstrate that it governs a pathogenic axis distinct from that of neuronal Lag3. We show that α-syn PFFs bind microglial Lag3 with high specificity and affinity, and that Lag3 is required for efficient uptake of α-syn PFFs into microglia. Microglial Lag3 deletion markedly suppresses microglial inflammatory activation, prevents the cytokine-driven induction of activated neurotoxic A1 astrocytes, and abolishes the astrocyte-dependent neuronal death triggered by α-syn PFF-activated microglia. Using microglia-specific Lag3 conditional knockout mice (Lag3^L/L^-Cx3cr1^CreER^), we further demonstrate that loss of microglial Lag3 attenuates pathologic α-syn neurodegeneration, reduces neuroinflammation, preserves dopaminergic neurons, and ameliorates motor deficits in vivo.

Together with our previous work on neuronal Lag3 [25], these findings support a unified, cell-type–specific model in which Lag3 contributes to α-syn pathology through two mechanistically distinct yet complementary pathways: neuronal Lag3 mediates α-syn PFFs binding, uptake, and trans-synaptic propagation, whereas microglial Lag3 recognizes extracellular pathologic α-syn PFFs to initiate inflammatory cascades that drive A1 astrocyte conversion and neurodegeneration. This dual-axis framework positions microglial Lag3 as a promising therapeutic target for mitigating pathologic α-syn-induced neuroinflammation and neurodegeneration in PD.

## Materials and Methods

### α-Synuclein Purification and Preparation of Preformed Fibrils (PFFs)

Recombinant mouse α-synuclein was expressed using the IPTG-independent pRK172 vector system and purified as previously described. Endotoxin was removed with the ToxinEraser endotoxin removal kit (Genscript). To generate α-synuclein preformed fibrils (PFFs), purified α-synuclein (5 mg/ml in PBS) was incubated at 37°C under continuous agitation (1,000 rpm) for 7 days. The resulting fibrillar aggregates were diluted to 0.1 mg/ml in PBS and sonicated for 30 s (0.5s on/off pulses) at 10% amplitude using a Branson Digital Sonifier (Danbury, CT, USA). PFFs formation was confirmed by atomic force microscopy (AFM), transmission electron microscopy (TEM), and by their capacity to induce pS129 α-syn immunoreactivity. Aliquots were stored at −80°C until use.

### Primary neuron, microglia, and astrocyte cell cultures

C57BL/6 and CD1 mice were obtained from Jackson Laboratories (Bar Harbor, ME). Primary cortical neurons were isolated from E15.5 embryos and cultured on poly-L-lysine–coated plates in Neurobasal medium (Gibco) supplemented with B-27, 0.5 mM L-glutamine, and penicillin-streptomycin (Invitrogen). Half of the culture medium was replaced every 3-4 days. Primary microglia and astrocytes were prepared from whole brains of postnatal day 1 (P1) pups as previously described [34]. Briefly, meninges were removed, and brains were washed three times in DMEM/F12 (Gibco) containing 10% heat-inactivated FBS, 50 U/mL penicillin, 50 μg/mL streptomycin, 2 mM L-glutamine, 100 μM non-essential amino acids, and 2 mM sodium pyruvate. Tissue was dissociated in 0.25% trypsin-EDTA for 10 min with gentle agitation, then quenched with complete DMEM/F12, and washed three additional times. The suspension was triturated, filtered through a 100-μm nylon mesh, and plated into T-75 flasks. Mixed glial cultures were maintained for 13 days, with a complete medium change on day 6.

Microglia-enriched and astrocyte-enriched fractions were obtained from mixed glial cultures using the EasySep Mouse CD11b Positive Selection Kit (StemCell). CD11b-positive microglia and CD11b-negative astrocytes were maintained separately. Conditioned medium from microglia treated with α-syn PFFs (α-syn PFFs-MCM) or PBS was collected and applied to primary astrocytes for 24 h. Astrocyte-conditioned medium activated by α-syn PFF-MCM (α-syn PFFs-ACM) was harvested, supplemented with Mini EDTA-free protease inhibitor cocktail (Sigma), and concentrated ∼50-fold using Amicon Ultra-15 centrifugal filters (10-kDa cutoff; Millipore). Protein concentration was quantified with the Pierce BCA assay (Thermo Scientific). For neuronal cell-death assays, 15 μg/mL of total α-syn PFF-ACM protein was added to mouse primary neurons.

### Microglia Cell Surface Binding Assays

Primary microglia isolated from WT and Lag3^−/−^ mice were incubated with α-syn–biotin PFFs (1 μM total α-syn-biotin monomer equivalent) in DMEM supplemented with 10% FBS for 2 h at room temperature. Following incubation, unbound material was removed by extensive washing (4-6 washes, 20 min each) with DMEM containing 10% FBS, and cells were subsequently fixed in 4% paraformaldehyde in PBS. Fixed microglia were rinsed with PBS and blocked for 30 min in PBS containing 10% horse serum and 0.1% Triton X-100. Cells were then incubated for 16 h at 4 °C with alkaline phosphatase–conjugated streptavidin (1:2000) diluted in PBS supplemented with 5% horse serum and 0.05% Triton X-100. Bound alkaline phosphatase was visualized using a BCIP/NBT chromogenic substrate. Quantification of α-syn-biotin PFFs binding was performed in ImageJ by applying a fixed threshold to define signal intensity and exclude background; identical threshold parameters were applied to all images within each experiment. Binding curves and dissociation constants (KD) were derived using nonlinear regression in GraphPad Prism.

### Live images

Primary microglia were plated on Nunc™ glass-bottom dishes for live imaging. α-syn PFFs were labeled with pHrodo Red (Invitrogen, Grand Island, NY, USA), a pH-sensitive fluorophore whose signal increases upon internalization into acidic microglial compartments. pHrodo-labeled α-syn PFFs were added directly to Lag3 WT and Lag3^−/−^ microglia, and live cell imaging was performed on a Zeiss Axio Observer Z1 microscope with images acquired every 0.5-1 min for 30-60 min. Cells became suitable for quantification approximately 1-2 min after exposure to α-syn PFFs. Fluorescence intensity was quantified by outlining individual microglia using Zeiss Zen software and subtracting background fluorescence. Baseline intensity was defined as the 2-3 min post-treatment signal, and the percentage of internalized α-syn-pHrodo at each time point was calculated relative to this baseline.

### Gene Expression and Cytokine Quantification

Total RNA from WT and Lag3^−/−^ primary microglia was isolated using the RNeasy RNA Isolation Kit (Qiagen, CA) according to the manufacturer’s instructions, and RNA quantity and purity were measured with a NanoDrop 2000 spectrophotometer (Biotek, Winooski, VT). For cDNA synthesis, 1–2 μg of total RNA was reverse-transcribed using the High-Capacity cDNA Reverse Transcription Kit (Life Technologies, Grand Island, NY). qRT-PCR was performed using custom TaqMan primer/probe sets and TaqMan Fast Advanced Master Mix on a QuantStudio 3 Real-Time PCR System. Before reverse transcription, samples were spiked with the TaqMan Universal RNA Spike-In RT Control (Thermo Fisher) as an internal reference. Gene expression was normalized to 18S rRNA and quantified using the comparative cycle threshold *Ct* method (2^−ΔΔ*Ct*^). Primer and probe sequences are listed in Table S1.

To assess cytokine secretion, culture supernatants collected 18 hours after α-syn PFFs treatment were processed using the Proteome Profiler Mouse XL Cytokine Array (R&D Systems, Minneapolis, USA) according to the manufacturer’s protocol.

### Propidium iodide staining

Primary cortical neurons were incubated for 24 h with α-syn PFFs-MCM-ACM derived from WT or Lag3⁻/⁻ microglia. Neuronal viability was assessed using unbiased, computer-assisted quantification following nuclear staining with 7 μM Hoechst 33342 to label all cells and 2 μM propidium iodide to identify non-viable nuclei. Total and dead cell numbers were quantified using Axiovision 4.3 software (Carl Zeiss). Percent cell death was calculated as the proportion of PI-positive cells normalized to control wells.

### Mouse Strain

Lag3^L/L-YFP^ mice were obtained from Dr. Dario Vignali (University of Pittsburgh), and Cx3cr1^CreER^ knock-in/knock-out mice were purchased from the Jackson Laboratory (strain #021160). Microglia-specific Lag3 conditional knockout mice (Lag3^L/L^-Cx3cr1^CreER^) were generated by crossing Lag3^L/L-YFP^ mice with Cx3cr1^CreER^ mice. Lag3^L/L^-Cx3cr1^CreER^ mice and littermate controls received intraperitoneal injections of tamoxifen (1.5 mg per mouse, once daily for five consecutive days) to induce Cre recombinase activity. In Cx3cr1^CreER^ mice, Cre recombinase is expressed in microglia as well as in peripheral myeloid populations. However, circulating myeloid cells undergo continuous turnover and are replaced within approximately one month by Cx3cr1-negative progenitors, resulting in loss of Cre activity in the periphery. Because microglia exhibit minimal turnover, tamoxifen-induced Cre expression is stably retained in these cells. Thus, a one-month washout period enables microglia-specific recombination.

C57BL/6 WT mice (strain #000664) and CD1 mice were obtained from the Jackson Laboratory (Bar Harbor, ME). Global Lag3^-/-^ mice were kindly provided by Dr. Charles G. Drake (formerly at Johns Hopkins University). All mice were maintained in individually ventilated cages with ad libitum access to food and water. All experimental procedures were approved by the Johns Hopkins Animal Care and Use Committee and conducted in accordance with NIH guidelines for the care and use of laboratory animals.

### Stereotaxic α-syn PFFs injection

For stereotaxic injection of α-syn PFFs, three-month-old male and female mice were anesthetized with ketamine and xylazine and secured in a stereotaxic frame. A 26.5-gauge injection cannula was positioned unilaterally into the right striatum using the following coordinates relative to bregma: mediolateral +2.0 mm, anteroposterior +0.2 mm, and dorsoventral −2.6 mm. α-syn PFFs (2 μL; 2.5 μg/μL in PBS) or an equal volume of PBS were infused at a rate of 0.2 μL/min. After completion of the infusion, the cannula was kept in place for 5 min before withdrawal to prevent backflow.

### Tissue Lysate Preparation and Immunoblot Analysis

Dissected brain regions were homogenized in RIPA lysis buffer supplemented with Protease/Phosphatase Inhibitor Cocktail (5872S; Cell Signaling Technology, Danvers, MA). Homogenates were centrifuged at 10,000 rpm for 20 min at 4°C, and the resulting supernatants were collected. Protein concentrations were quantified using the Pierce BCA assay (Thermo Scientific, USA). Equal amounts of protein (20–30 µg) were separated on 12.5–13.5% SDS-polyacrylamide gels and transferred to PVDF membranes. Membranes were blocked with 5% nonfat milk or 5% BSA in TBST (0.1% Tween-20), incubated with primary and HRP-conjugated secondary antibodies, and developed using ECL or SuperSignal Femto substrates (Thermo Fisher, USA). Chemiluminescent signals were detected using an ImageQuant LAS 4000 mini system (GE Healthcare Life Sciences). All primary and secondary antibodies used for immunoblotting are listed in Table S2.

### Immunohistochemistry and Stereological Quantification

Immunohistochemistry was performed on free-floating 30-µm coronal brain sections. Primary antibodies and working dilutions are listed in the Supplementary Table 2. Staining and quantification of TH-, Nissl-positive neurons were conducted according to established protocols in which the examiner was blinded to the experimental groups. Microglia within the substantia nigra pars compacta (SNpc) were labeled using anti-Iba1 (Wako, St. Louis, MO, USA) antibodies followed by biotin-conjugated anti-rabbit secondary antibodies and ABC reagents. Sections were developed using SigmaFast DAB Peroxidase Substrate (Sigma-Aldrich). Microglial densities in the SNpc were quantified using ImageJ.

### Immunofluorescence Analysis

Immunofluorescence staining was performed on free-floating 30 µm brain sections, using primary antibodies and working dilutions listed in the Supplementary Table 2. Tissue immunofluorescence procedures and analysis followed established methods [23, 25]}.

For cultured cell staining, paraformaldehyde-fixed cells were rinsed once with PBS, then blocked and permeabilized with 10% normal goat serum (NGS; Vector Labs) containing 0.1% Triton X-100 in PBS for 15–30 min at room temperature. Primary antibodies were applied for 1 hour at room temperature or overnight at 4° C in 10% NGS/PBS. After two PBS washes, Alexa Fluor–conjugated secondary antibodies (Thermo Fisher) were applied for 1 h at room temperature in 5% NGS/PBS. Cells were rinsed in PBS containing Hoechst 33342. Imaging of tissue sections and cultured cells was performed on a laser-scanning confocal microscope (LSM 880, Zeiss, Dublin, CA, USA) or a fluorescent microscope (BZ-X710, Keyence, Osaka, Japan).

### Behavioral Assessments

Behavioral testing was performed 10 days before euthanasia to assess motor deficits induced by α-syn PFFs or PBS. All assays were conducted by investigators blinded to treatment. The behavioral battery included the grip strength test, pole test, rotarod test, and wire hanging test.

Grip strength was assessed using a Bioseb force transducer. Mice grasped a metal grid with their forelimbs or all four limbs, and the peak force before grip release was recorded in grams. For the pole test, mice were acclimated for 30 min and trained for two days (three trials/day). On the test day, mice were placed 7.5 cm from the top of a 75-cm gauze-wrapped vertical pole (9 mm in diameter), and the time required to turn downward and descend to the base was recorded (three trials; 60-s cutoff). Turn-down time, climb-down time, and total descent time were quantified.

For the rotarod test, mice were trained for three days before testing. On the test day, they were placed on an accelerating rotarod (4-40 rpm over 5 min), and latency to fall was recorded in three trials. Trials were terminated after two passive complete rotations. Performance was reported as the mean latency to fall relative to controls.

In the wire-hanging test, mice were placed on a 2 mm horizontal wire suspended 30 cm above the bedding. Latency to fall (maximum 60 s) and posture-based grip performance were recorded across three trials to assess muscle endurance and grip strength.

## Results

Lag3 was detected in both neurons and microglia, with stronger expression in microglia. To examine whether extracellular α-syn PFFs interact with microglia, we used biotinylated α-syn PFFs. Streptavidin-AP staining showed that α-syn-biotin PFFs robustly bound to microglia, whereas α-syn-biotin monomers exhibited only weak, nonspecific interactions (Fig. 1A, B; Fig. S1A). Binding of α-syn-biotin PFF to microglia was saturable, with an apparent dissociation constant (Kd) of 92.2 nM (Fig. 1A, B; Fig. S1A). Consistently, WT microglia showed strong α-syn-biotin PFFs binding, while Lag3^−/−^ microglia displayed markedly reduced binding. By subtracting binding in Lag3^−/−^ cultures from that in WT cultures, we identified specific Lag3-dependent binding with a Kd of 5.6 nM (Fig. 1A, B). These findings indicate that α-syn PFFs bind extracellular Lag3 on microglia with high specificity.

**Fig. 1.**
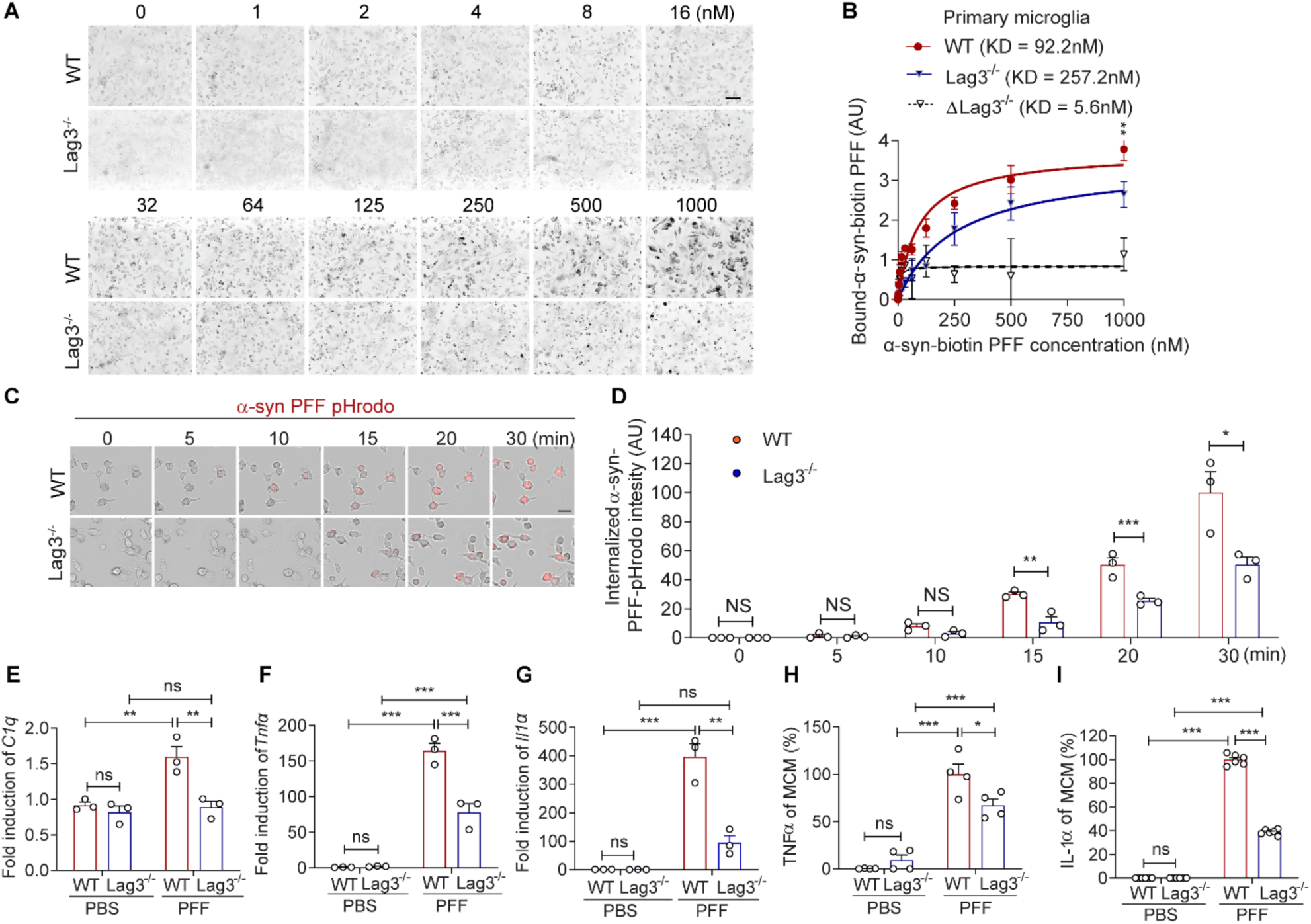
Inhibition of α-syn PFFs-induced microglial activation by Lag3^-/-^. **(A-B)**, Comparison of α-syn-biotin PFFs binding to wild-type (WT) and Lag3^-/-^ mouse primary microglia (dotted line, ΔLag3 = wild type minus Lag3^−/−^). **(C-D)**, Live image and quantification of the intensity of α-syn-pHrodo PFFs in WT and Lag3^-/-^ mouse primary microglia from 3 independent experiments. Two-way ANOVA with Tukey’s multiple comparisons test. Scale bar, 50μm. **(E-G)**, qPCR analysis of α-syn PFFs-induced microglial activation markers. (**E)** *C1q* expression, (**F**) *Tnfα* expression, (**G**) *Il1α* expression. Data means s.e.m.; n=3 biologically independent primary microglia cultures. **(H-I)**, Cytokine analysis of α-syn PFFs-activated microglia-conditioned medium (MCM) 18h after α-syn PFF treatment by ELISA. The increase of TNFα (**H**) and IL-1α (**I**) is prevented by Lag3^-/-^. Data: mean ± SEM; n=3 sample treated with primary microglia-conditioned medium. \**P* < 0.05; \*\**P* < 0.01; \*\*\**P* < 0.001. ns = no significance.

We next asked whether Lag3 also contributes to microglial α-syn PFFs uptake. To test this, α-syn PFFs were conjugated to pHrodo red, a pH-sensitive dye that fluoresces more intensely in the acidic environment of endosomes than in the neutral cytosol. In WT microglial cultures, α-syn-pHrodo PFFs were efficiently internalized, as evidenced by a time-dependent increase in fluorescence. By contrast, Lag3^−/−^ microglia exhibited minimal uptake under the same conditions (Fig. 1C, D). Together, these results suggest that Lag3 not only mediates extracellular binding of α-syn PFFs but also facilitates their subsequent endocytosis in microglia.

To determine whether depletion of microglial Lag3 alters the release of factors that drive neurotoxic reactive astrocyte conversion, astrocytes were treated with MCM collected from WT or Lag3^−/−^ microglia exposed to α-syn PFFs for 24 h (Fig. 2A). After incubation, astrocytic C3 expression, a hallmark of A1 astrocyte activation, was assessed. α-syn PFFs-MCM strongly induced C3 expression in astrocytes, whereas this induction was markedly suppressed when astrocytes were treated with MCM from Lag3^−/−^ microglia (Fig. 2. B, C). To further define the astrocytic phenotype, we quantified general reactive, A1-, and A2-specific transcripts by qPCR (Fig. 2D-F). α-syn PFFs-MCM preferentially upregulated A1-specific markers, while leaving most A2 transcripts unchanged, except for Ptx3. These results indicate that α-syn PFFs-activated microglia specifically convert astrocytes into neurotoxic reactive A1 astrocytes, and that depletion of microglial Lag3 effectively prevents this A1-specific induction. Consistent with the primary microglia and astrocytes results, intrastriatal injection of α-syn PFFs into Lag3^L/L^ and Lag3^L/L^-Cx3cr1^CreER^ mice induces C3/GFAP expression in the cortex, but the C3/GFAP expression is significantly reduced in the Lag3^L/L^-Cx3cr1^CreER^ mice (Fig. 3H-J).

**Fig 2.**
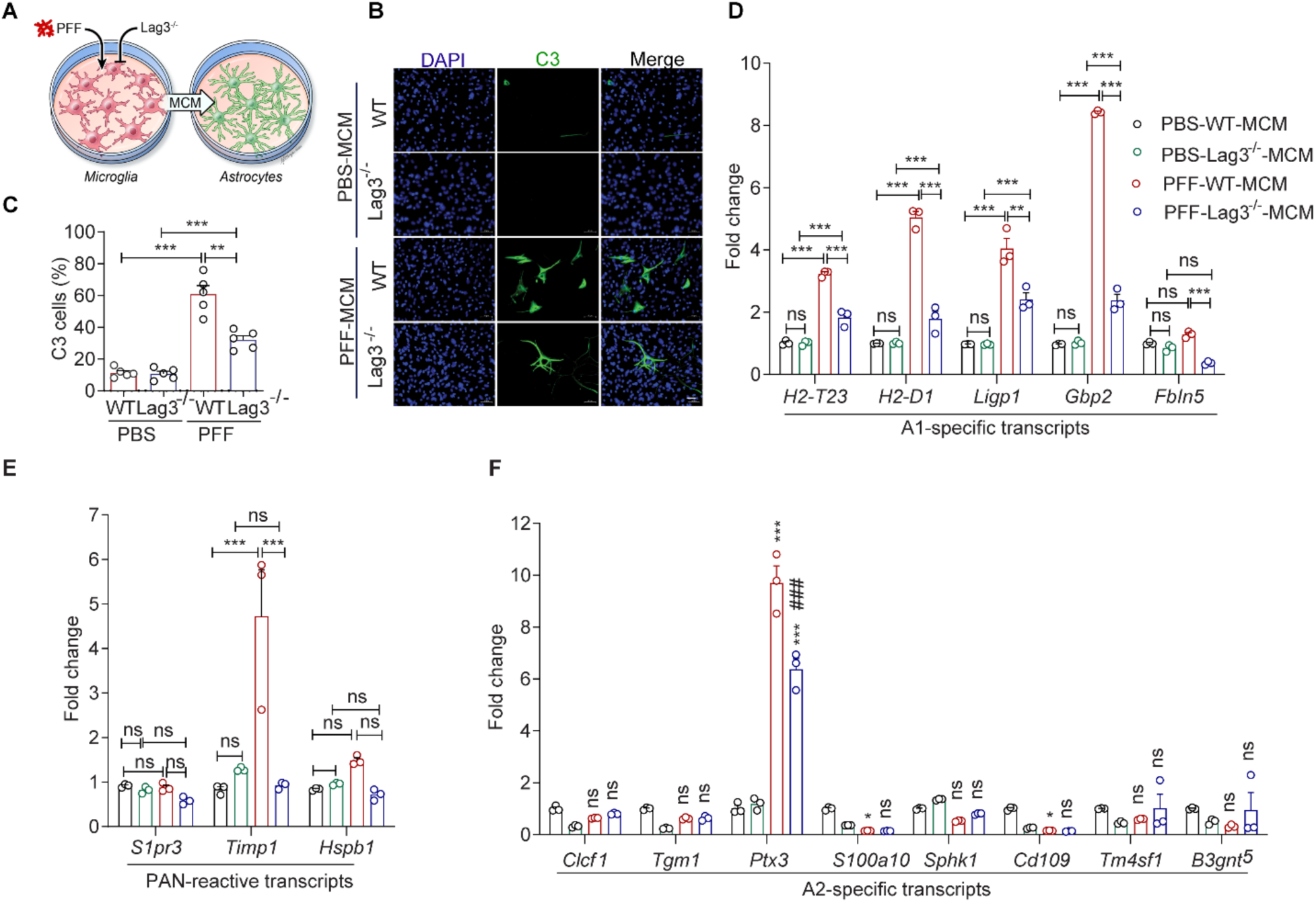
Inhibition of α-syn PFF-induced neurotoxic reactive A1 astrocytes and related neurotoxicity by Lag3^-/-^. **(A-C)**, C3 expression in the primary culture of astrocytes was assessed by confocal microscopy. Scale bar, 10 μm. Astrocytes treated with α-syn PFFs-microglia conditional medium (MCM) from WT and Lag3^-/-^ microglia. Data means s.e.m.; n=3 biologically independent primary astrocyte and microglia cultures. **(D-F)**, Inhibition of α-syn PFFs-MCM induced A1 reactive astrocytes by Lag3^-/-^ in purified primary astrocytes. Fold change in mRNA expression of the indicated genes. Data: mean ± SEM; n=3 biologically independent primary astrocyte cultures. \*\**P* < 0.01; \*\*\**P* < 0.001. ns = no significance.

**Fig 3.**
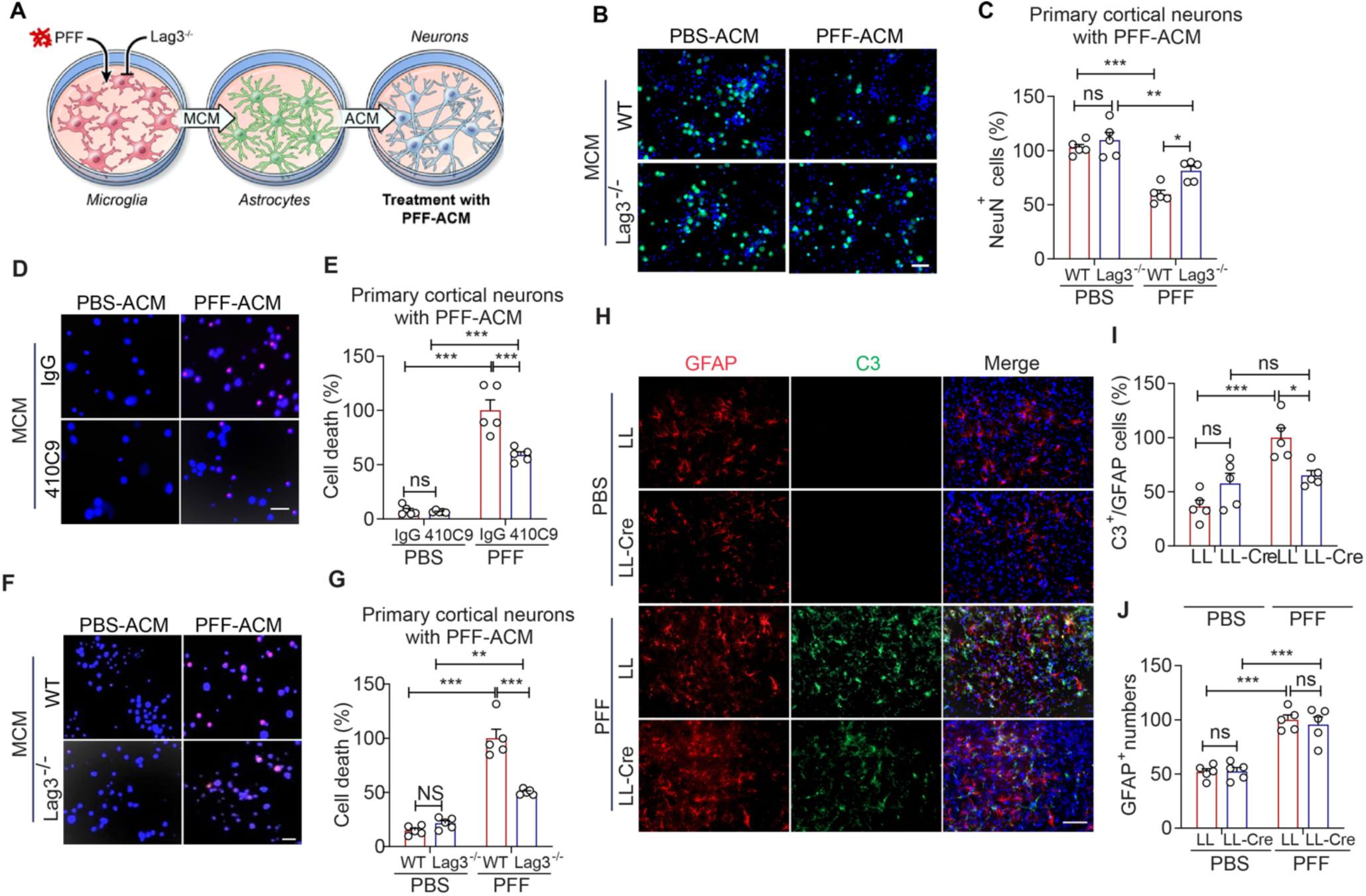
**(A-C)** Lag3^-/-^ prevents α-syn PFFs-MCM-ACM toxicity as assessed by NeuN staining. (**D-E)** Lag3 antibody prevents α-syn PFFs-MCM-ACM toxicity as assessed by Hoechst and PI staining. (**F-G)** Lag3^-/-^ prevents α-syn PFFs-MCM-ACM toxicity as assessed by Hoechst and PI staining. Data: mean ± SEM; . n=3 biologically independent primary cortical neurons. Scale bar, 20 μm. (**H-J**) Co-localization of C3 and GFAP as assessed by confocal microscopy in vivo. Scale bar, 100 μm. **I,** Quantification of increased C3 transcripts, which is prevented in Lag3^L/L^-Cx3cr1^CreER^ mice injected with α-syn PFF. Data: mean ± SEM; n=5 biologically independent animals. \**P* < 0.05; \*\**P* < 0.01; \*\*\**P* < 0.001. ns = no significance.

We next investigated whether astrocytic conversion influences neuronal viability. Astrocytes were incubated with MCM for 24 h, after which ACM was collected, concentrated, and applied to primary mouse cortical neuron cultures (Fig. 3A). ACM derived from α-syn PFFs-treated WT microglia induced significant neuronal death, as assessed by NeuN and PI staining. In contrast, ACM generated following exposure to Lag3⁻/⁻ microglia failed to elicit comparable neurotoxicity and significantly attenuated α-syn PFFs-MCM-ACM-induced neuronal cell death (Fig. 3B-G). Collectively, these results indicate that microglial Lag3 deficiency confers neuroprotection, likely by limiting the release of proinflammatory factors that drive astrocytic conversion toward a neurotoxic A1 phenotype.

The potential of conditional Lag3 deletion in microglia on pathologic α-syn induced degeneration was evaluated in the stereotactic intrastriatal α-syn PFFs mouse model of sporadic Parkinson’s disease. α-syn PFFs were injected intrastriatally into Lag3^L/L^ and Lag3^L/L^-Cx3cr1^CreER^ mice (Fig. 4A). At 3 months post-injection, immunostaining confirmed robust induction of pS129 α-syn immunoreactivity in the cortex, striatum, and substantia nigra of α-syn PFF-injected mice, consistent with widespread α-syn pathology. Strikingly, Lag3^L/L^-Cx3cr1^CreER^ mice showed a significant decrease in pS129 α-syn immunoreactivity in these regions (Fig. 4B, C). At 6 months, α-syn PFFs injection induced a significant loss of tyrosine hydroxylase (TH) and Nissl-positive neurons in the substantia nigra pars compacta (SNpc), which was prevented in Lag3^L/L^-Cx3cr1^CreER^ mice (Fig. 3D-F). Western blot analysis further revealed that the α-syn PFFs-mediated reductions in TH and dopamine transporter (DAT) expression in the ventral midbrain were restored in Lag3^L/L^-Cx3cr1^CreER^ mice (Fig. 4G-I). We next evaluated motor function using multiple behavioral assays at 6 months post-injection. As expected, α-syn PFF-injected mice exhibited significantly reduced grip strength in both forelimbs and all four limbs, along with impaired performance in the accelerating pole test, rotarod test, and wire hanging tests. Notably, Lag3^L/L^-Cx3cr1^CreER^ mice were protected from these deficits, showing significantly improved performance across all behavioral measures (Fig. 4J–N). Finally, to determine whether these protective effects were associated with suppression of microglial activation, we examined IBA1 immunoreactivity. Intrastriatal injection of α-syn PFFs significantly increased IBA1 staining and microglial density in the SNpc in 6 months, whereas Lag3^L/L^-Cx3cr1^CreER^ mice exhibited markedly reduced IBA1 immunoreactivity (Fig. S2A, B). Western blot analysis of IBA1 confirmed these findings, demonstrating α-syn PFFs-induced microglial activation in WT mice and its attenuation in Lag3^L/L^-Cx3cr1^CreER^ mice (Fig. S2C, D).

**Fig 4.**
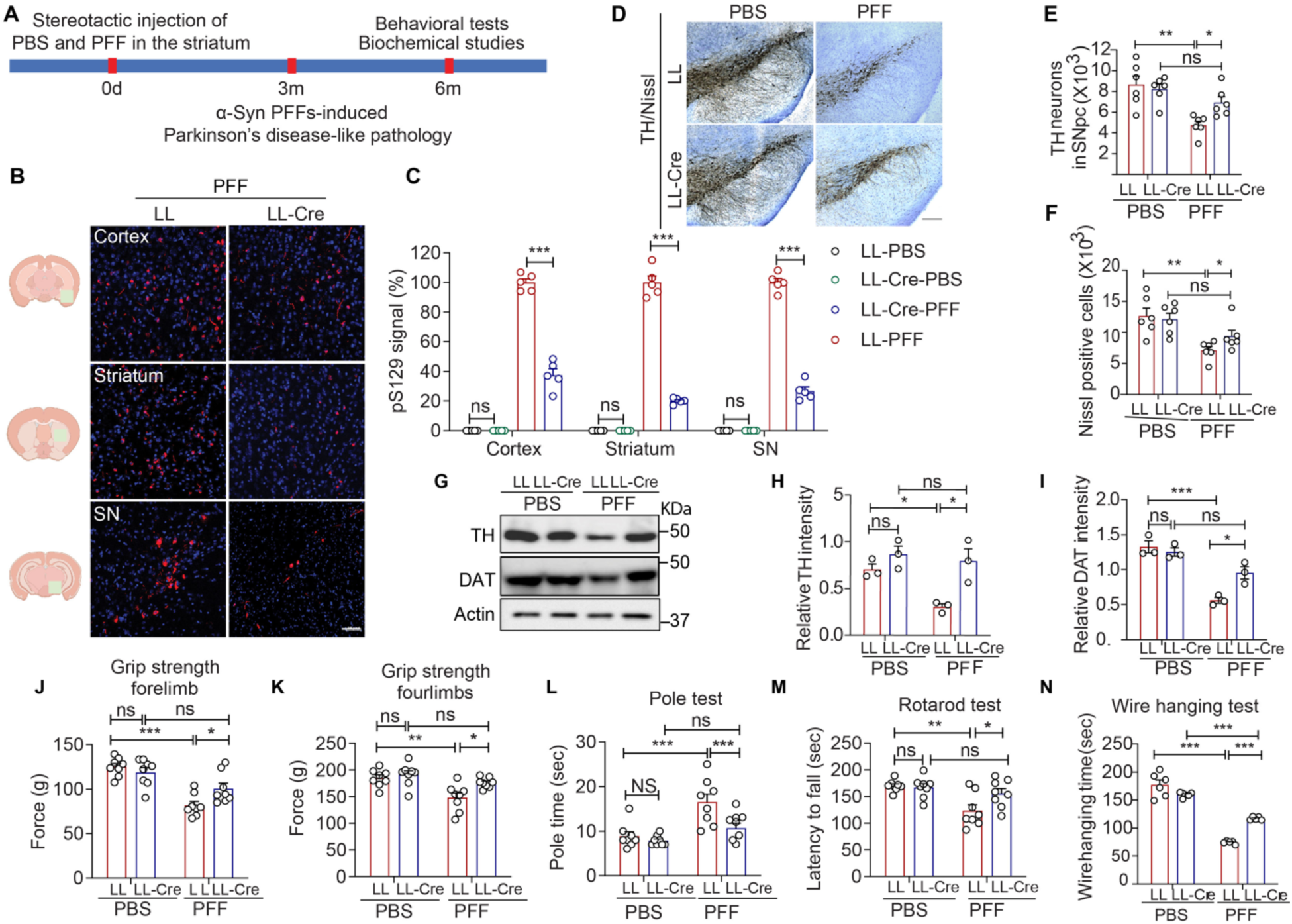
**(A)** Schematic diagram of the α-syn PFF experimental design. (**B-C**) Representative immunostaining for pS129 α-syn in the Cortex, Striatum, and SNpc. Green box in the left panel of the outlines area. Scale bars, 100 μm. (**D**) Representative photomicrographs from coronal mesencephalon sections containing tyrosine hydroxylase-positive neurons in the SNpc region. n=6 biologically independent animals. Scale bar, 500 μm. Unbiased stereological counts of (**E)** tyrosine hydroxylase-positive and (**F**) Nissl-positive neurons in the SNpc region. Data: mean ± SEM; n=6, biologically independent animals. unpaired two-tailed Student’s T-tests were used for statistical analyses. \*\*\**P* <0.001, NS, not significant. (**G-I)** Lag3^L/L^-Cx3cr1^CreER^ rescues PFF reduction in TH and DAT levels in the ventral midbrain. (**G**) Representative immunoblots of TH, DAT, and β-actin in the ventral midbrain. (**H-I**) Quantification of TH and DAT protein levels normalized to β-actin. Data: mean ± SEM; n=3, biologically independent animals. Ordinary one-way ANOVA with Tukey’s multiple comparisons test. (**J-N)**, Behavioral tests after PBS or α-syn PFF stereotaxic intrastriatal injection at six months in vivo. (**J-K**) grip strength of the forelimb and four limbs. (**L**) pole test. The maximum time allowed to climb down the pole was 60 seconds. (**M**) rotarod (**N**) wire hanging test. Data: mean ± SEM; n = 6∼10 biologically independent animals. A two-way ANOVA was used for statistical analysis, followed by a Tukey’s multiple comparisons test. \**P* < 0.05, \*\**P* < 0.01, \*\*\**P* < 0.001, ns = no significance.

Together, these results demonstrate that conditional deletion of Lag3 in microglia attenuates α-syn PFFs pathology, preserves dopaminergic neurons, restores striatal dopaminergic markers, improves motor performance, and suppresses microglial activation in vivo.

## Discussion

In this study, microglial Lag3 was identified as a key receptor for pathologic α-syn, and it was shown to drive an inflammatory cascade culminating in neurotoxic reactive A1 astrocyte conversion and neurodegeneration. These findings broaden the functional landscape of Lag3 in α-synucleinopathies and expand the cellular context in which Lag3 contributes to disease progression.

Our findings demonstrate that α-syn PFFs bind microglial Lag3 with high specificity, and Lag3 is required for efficient fibril uptake and microglial activation. Loss of Lag3 in microglia markedly reduces cytokine production and prevents the release of factors that induce neurotoxic A1 astrocytes, positioning microglial Lag3 as an upstream regulator of a glia-driven pathogenic cascade. In vivo, microglia-specific deletion of Lag3 attenuates pS129 α-syn pathology, suppresses microgliosis, limits A1 astrocyte induction, preserves nigrostriatal dopaminergic neurons, and improves motor performance following α-syn PFFs inoculation. These findings establish microglial Lag3 as an important mediator linking pathologic α-syn recognition to inflammatory amplification and neurodegeneration [35]. Beyond establishing microglial Lag3 as a critical node in α-syn PFF-induced neuroinflammation, our work highlights the role of microglial Lag3 in sensing pathologic α-syn and its role in microglia activation and conversion of neurotoxic reactive A1 astrocytes.

Work in multiple systems, including human iPSC-derived dopaminergic neurons, shows that even low-abundance neuronal Lag3 can mediate α-syn PFF binding, uptake, and pathogenic spread [11, 12, 23–25]. These findings support a cell-type- and context-dependent model in which neuronal Lag3 promotes pathologic α-syn transmission, while microglial Lag3 coordinates innate immune sensing and inflammatory amplification.

This dual-axis framework unites two distinct mechanisms: a neuronal propagation axis that drives α-syn PFFs dissemination, and a microglial neuroinflammation axis that induces A1 astrocyte conversion and consequent neurotoxicity. Viewing Lag3 function through this cell-type-specific lens emphasizes the need to study Lag3 within defined cellular compartments. Moreover, the identification of microglial Lag3 as an upstream regulator mechanistically links pathologic α-syn to glial network dysfunction, positioning Lag3 at the intersection of protein aggregation and neuroinflammation.

Our results also have important therapeutic implications. Selective inhibition of Lag3 in microglia may attenuate α-syn-driven inflammation while minimizing systemic immunosuppression. Interventions that block Lag3 pathologic α-syn binding, such as brain-penetrant antibodies, nanobodies, aptamers, or microglia-targeted gene-silencing strategies, could simultaneously reduce pathologic α-syn uptake and disrupt downstream inflammatory cascades. Furthermore, combined targeting of both neuronal and microglial Lag3 may offer a more comprehensive therapeutic approach by concurrently limiting pathologic α-syn propagation and neuroinflammation [10, 36].

Several questions remain for future investigation, including the intracellular signaling pathways downstream of Lag3 engagement, potential crosstalk with established innate immune receptors, and the contribution of microglial Lag3 to human disease progression [27]. Understanding how Lag3 integrates with broader glial networks and determining the structural basis for high-affinity α-syn PFFs binding will further refine strategies to target this receptor.

In summary, this study establishes microglial Lag3 as a central regulator of α-syn PFFs recognition, microglial activation, astrocyte conversion, and neurodegeneration. Together with prior work on neuronal Lag3, these findings provide a unified, cell-type-specific framework for understanding Lag3’s multifaceted roles in α-synucleinopathies. The distinct yet complementary functions of Lag3 in neurons and microglia highlight its potential as a precision therapeutic target for modifying disease progression in PD.

## Acknowledgements

The authors acknowledge the joint participation by the Adrienne Helis Malvin Medical Research Foundation through its direct engagement in the continuous active conduct of medical research in conjunction with The Johns Hopkins Hospital and the Johns Hopkins University School of Medicine and the Foundation’s Parkinson’s Disease Program M-2023. TMD is the Leonard and Madlyn Abramson Professor in Neurodegenerative Diseases. This work was supported by the Freedom Together Foundation and the Robert J. and Claire Pasarow Foundation and NIH/NIA R01 AG085688 (VLD, TMD). T.M.D is the Leonard and Madlyn Abramson Professor in Neurodegenerative Diseases. The Multiphoton Imaging Core of Johns Hopkins University was used (NS050274) in some of the imaging studies. JTH received support through the Medical Scientist Training Program at Johns Hopkins University School of Medicine (NIH/NIGMS T32 GM136577) and through a predoctoral NRSA (NIH/NIA F30AG067643).

## Author Contributions

Conceptualization, X.M. V.L.D., T.M.D; Methodology, X.Y., R.K., J.T.H. ; Validation, X.Y., R.K., ; Formal Analysis, X.Y., R.K., ; Investigation, X.Y., R.K., ; Resources, T.M.D., V.L.D.; Writing-Original Draft, X.Y., R.K., T.M.D., V.L.D.; Visualization, X.Y., R.K., ; Supervision, X.M. T.M.D., V.L.D.; Funding Acquisition, X.M., T.M.D., V.L.D.

---

**Fig. S1.**
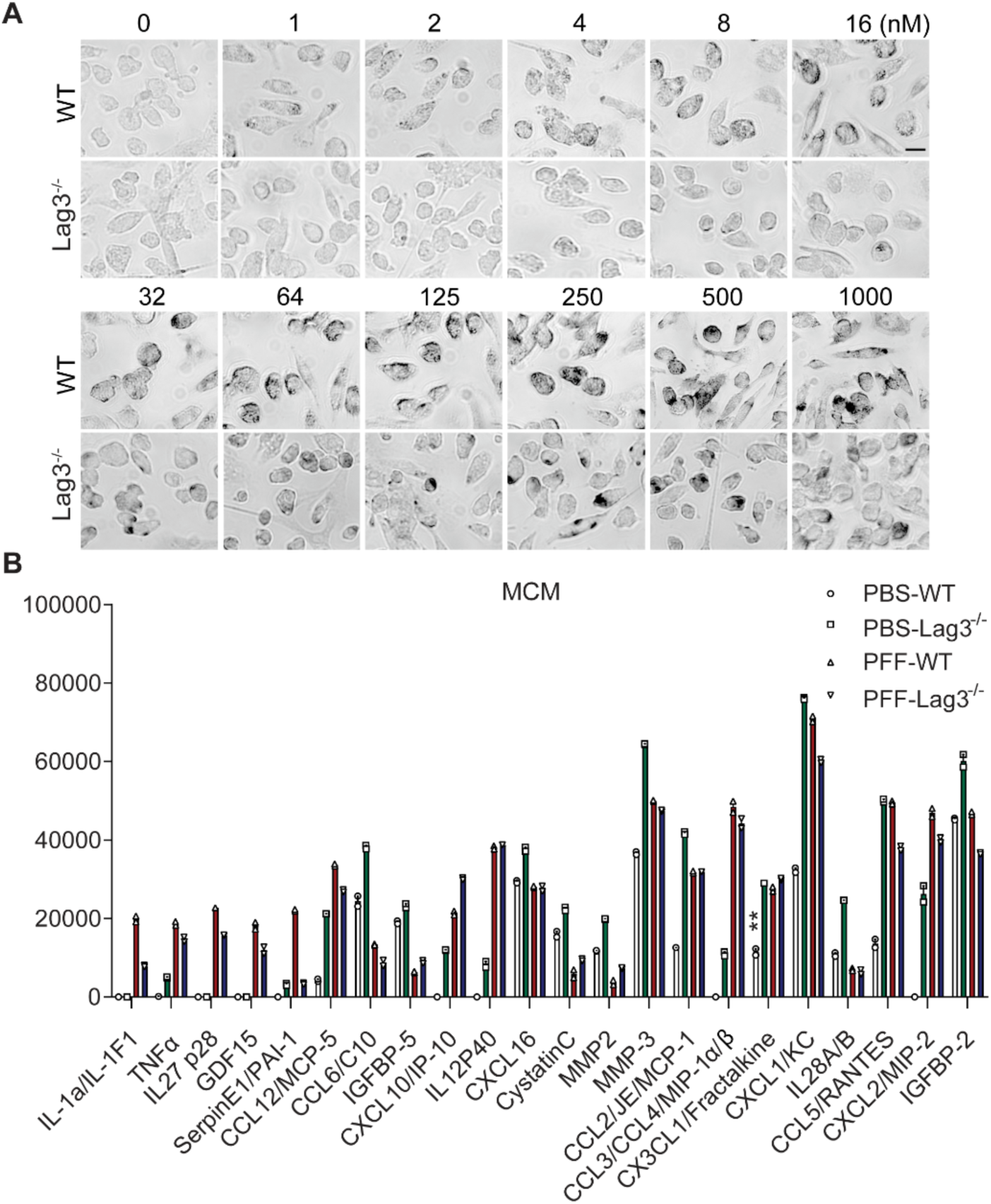
Characterization of α-syn-biotin PFFs binding to wild-type (WT) and Lag3^-/-^ mouse primary microglia. **(A)**, Comparison of α-syn-biotin PFFs binding to WT and Lag3^-/-^ mouse primary microglia. **(B)**, Cytokines in microglia-conditioned medium (MCM) collected 18 h after α-syn PFF treatment were quantified using ELISA.

**Fig. S2.**
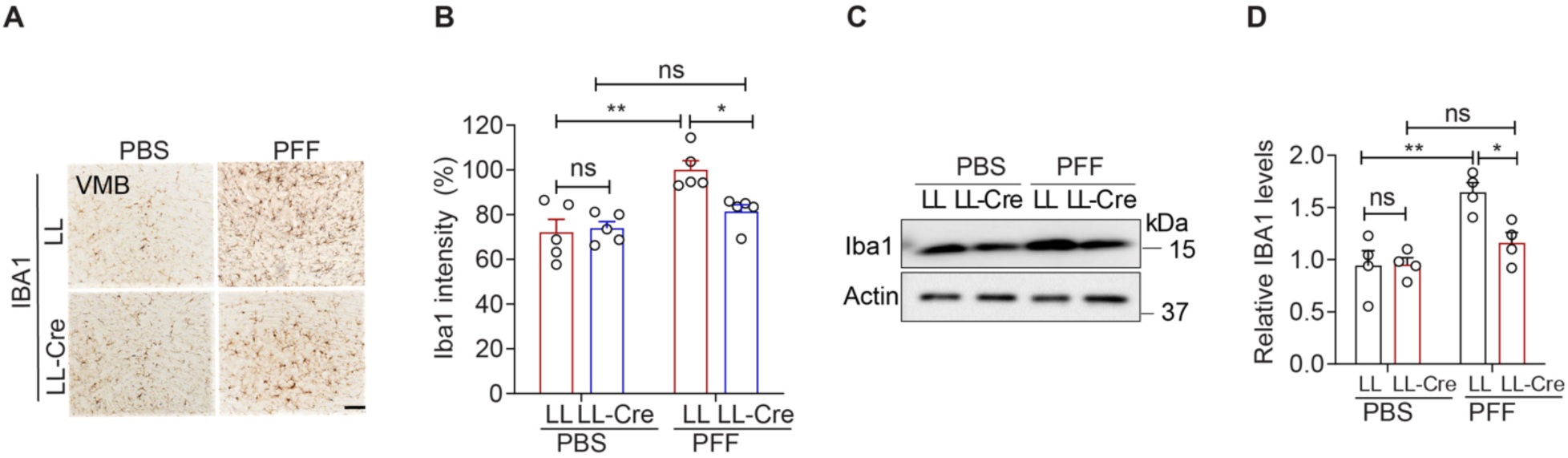
Inhibition of α-syn PFFs-induced microglial activation by Lag3^L/L^-Cx3cr1^CreER^. (A) Representative immunohistochemical images of IBA1. Scale bar, 100 μm. (B) Intensity of **I**BA1-positive signals in the SNpc. Data: mean ± SEM; n=5 biologically independent animals. (**C**) Representative immunoblots of IBA1 and β-actin in the ventral midbrain of PBS and α-syn PFF-injected mice. (**D**) Quantification of IBA1 levels in ventral midbrain normalized to β-actin. Data: mean ± SEM; n=3 biologically independent animals. One-way ANOVA followed by Tukey’s multiple comparisons test. \**P* < 0.05, \*\**P* < 0.01, ns = no significance.

**Table S1:**
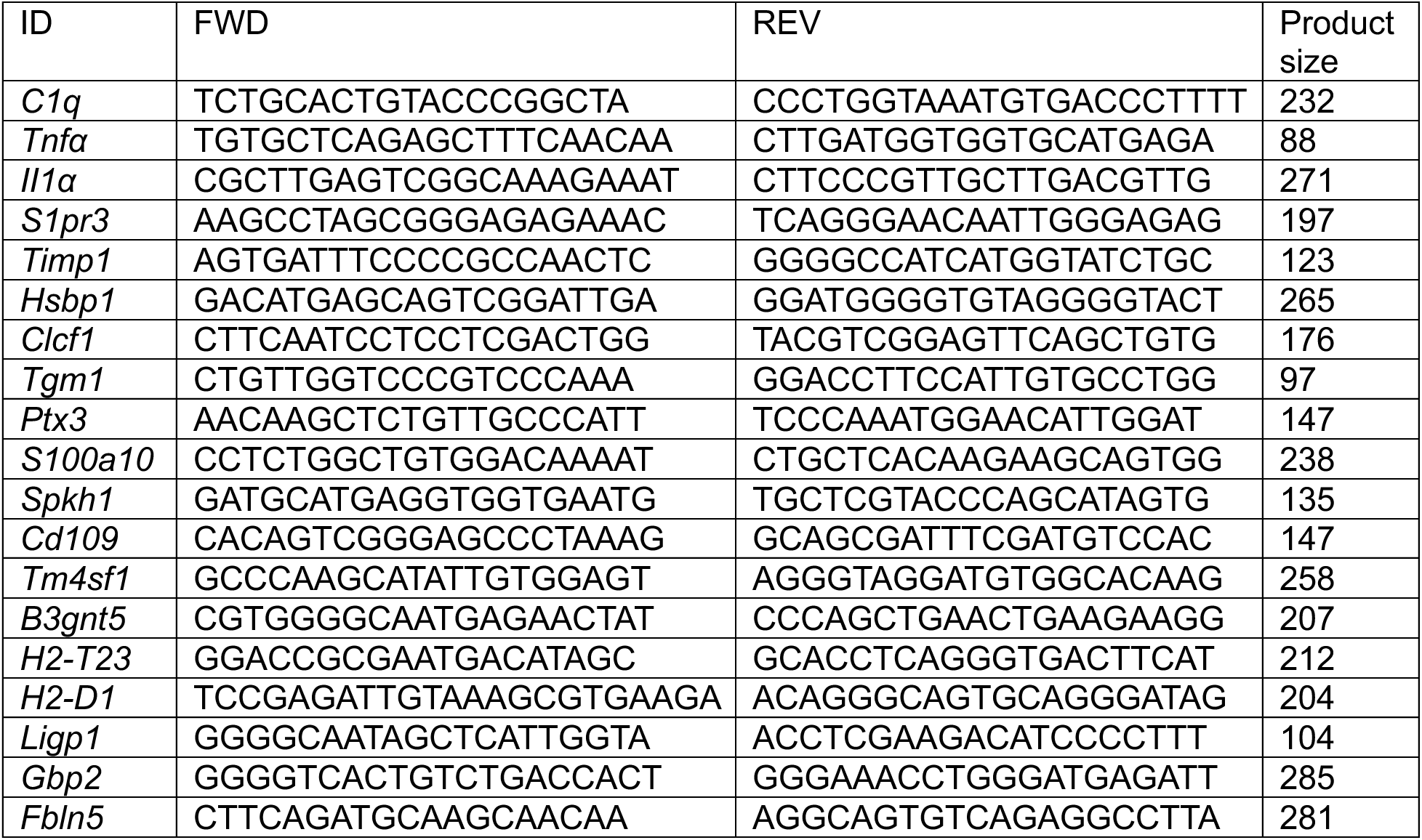
Mouse primer sequences.

**Table S2:** Antibodies.

| Antibodies | Source/Cat.No | Host | Dilution |
| --- | --- | --- | --- |
| p-α-synucleinSer129 | Abcam (ab51253) | Rabbit | 1:1,000 (IF) |
| Tyrosine Hydroxylase (TH) | Novus Biologicals (NB300-109) | Rabbit | 1:2,000 (WB)<br>1:1,000 (IHC, IF) |
| Dopamine transporter (DAT) | Sigma (D6944) | Rabbit | 1:1,000 (WB) |
| GFAP | Dako (Z0334) | Rabbit | 1:500 (IF) |
| Iba-1 | Wako (019-19741) | Rabbit | 1:500 (IHC)<br>1:1,000 (WB) |
| C3 (B-9) | Santa Cruz (sc-28294) | Mouse | 1:200 (IF) |
| β-actin-HRP | Sigma-Aldrich (A3854) | Mouse | 1:50,000 (WB) |
| Normal mouse IgG | Santa Cruz (sc-2025) | Mouse | 10 μg/ml (neutralizing) |
| NeuN | ab104224 | Mouse | 1:200 (IF) |
| Hoechst 33342 | Thermo Scientific (62249) |  | 1:2,000 (IF) |
| Biotin | Sigma (G3893) | Mouse | 1:1,000 (IF) |
| Goat Anti-Mouse IgG (H+L) Antibody, Alexa Fluor 488 Conjugated | Thermo Fisher Scientific (A11001) | Goat | 1:500 |
| Goat anti-Rabbit IgG (H+L) Cross-Adsorbed Secondary Antibody, Alexa Fluor™ 568 | Thermo Fisher Scientific (A11011) | Goat | 1:500 |
| ECL™ Anti-Rabbit IgG, Horseradish Peroxidase | Millipore (NA934V) | Donkey | 1:5,000 (WB) |
| Streptavidin, Alexa Fluor™ 647 Conjugate | Thermo Fisher Scientific (s-21347) |  | 1:1,000 (IF) |
| Protease/Phosphatase Inhibitor Cocktail | Cell Signaling Technology (5872S) |  | 1:100 (WB) |

